# Automatic generation of ground truth data for the evaluation of clonal grouping methods in B-cell populations

**DOI:** 10.1101/2020.11.30.404046

**Authors:** Nika Abdollahi, Anne de Septenville, Frédéric Davi, Juliana S. Bernardes

## Abstract

**Motivation:** The adaptive B-cell response is driven by the expansion, somatic hypermutation, and selection of B-cell clones. Their number, size and sequence diversity are essential characteristics of B-cell populations. Identifying clones in B-cell populations is central to several repertoire studies such as statistical analysis, repertoire comparisons, and clonal tracking. Several clonal grouping methods have been developed to group sequences from B-cell immune repertoires. Such methods have been principally evaluated on simulated benchmarks since experimental data containing clonally related sequences can be difficult to obtain. However, experimental data might contains multiple sources of sequence variability hampering their artificial reproduction. Therefore, the generation of high precision ground truth data that preserves real repertoire distributions is necessary to accurately evaluate clonal grouping methods.

**Results:** We proposed a novel methodology to generate ground truth data sets from real repertoires. Our procedure requires V(D)J annotations to obtain the initial clones, and iteratively apply an optimisation step that moves sequences among clones to increase their cohesion and separation. We first showed that our method was able to identify clonally-related sequences in simulated repertoires with higher mutation rates, accurately. Next, we demonstrated how real benchmarks (generated by our method) constitute a challenge for clonal grouping methods, when comparing the performance of a widely used clonal grouping algorithm on several generated benchmarks. Our method can be used to generate a high number of benchmarks and contribute to construct more accurate clonal grouping tools.

**Availability and implementation:** The source code and generated data sets are freely available at github.com/NikaAb/BCR_GTG

## 1 Introduction

B lymphocytes are a major cellular component of the adaptive immune system. They recognize and directly bind to pathogens (antigens) through a specific receptor, the B-cell receptor (BCR), located on the outer cell surface. The BCR consists of two components: the recognition unit, structured by a membrane immunoglobulin (IG) protein, and an associated signalling unit. Upon antigen-specific stimulation, B lymphocytes undergo activation, proliferation and further differentiation allowing the secretion of antibodies, a soluble form of the IG. Immunoglobulins are composed of two identical heavy chains (IGH) and two identical light chains (IGL). Each chain has two distinct parts: the variable region in the N-terminal side responsible for antigen recognition and the constant region on the C-terminal side anchored to the cell membrane. In this work, we use only IGH sequences, since they are more diverse than IGL chains, providing a more reliable signature for immune repertoire studies (*1*).

Complex genetic mechanisms create tremendous diversity of IGH variable regions. Three sets of genes encode these regions: variable (V), diversity (D), and joining (J). These genes are naturally separated on the genome. However, they become juxtaposed during early B-cell ontogeny by a process called VDJ recombination, which randomly selects and joins together one of each of three types of genes (*2*). Joining is imprecise as nucleotides are randomly deleted and inserted in the V-D (N1) and D-J (N2) junctions. The N1-D-N2 region is at the centre of the so-called third complementarity determining region (in short CDR3) and has the highest variability within the IGH molecule. VJ recombination also occurs for IGL genes, and random pairing of IGH and IGL chains allows the production of highly diverse B lymphocytes.

Upon antigen activation, B cells undergo rapid proliferation (clones) and further diversification of their BCR sequences by somatic hypermutation (SHM), an enzymatically-driven process introducing nucleotide substitutions into the IG variable genes. Consequently, each B lymphocyte expresses a unique IG nucleotide sequence, which enables the recognition of a particular set of antigens. The sum of all B cells with distinct BCRs is termed the BCR repertoire.

B-cell clones are essential cellular units of the immune system. They consist of groups of identical cells which derive from a common precursor. However, due to the complexity of the B-cell differentiation process, mature B cell clones, although sharing the same IGH and IGL rearrangements, can contain many molecular variants due to SHM. Identifying clones in BCR repertoires (clonal grouping) is the starting point for several studies involving distinct subjects like: autoimmune disease (*3*), allergy (*4*), cancer (*5*), ageing (*6*), and SARS-CoV-2 infection (*7*). Another important application of the clonal grouping is to distinguish between clonal (tumoral) or non-clonal (non-tumoral) B-cell populations in case of suspicion of B-cell malignancies (*8*). Extremely varied BCR repertoires are termed as polyclonal (or non-clonal) and are generally observed in healthy individuals. In contrast, malignant lymphoid cells (leukemias/lymphomas) consist of monoclonal populations. In-between these two extreme situations, the immune repertoire can display unique or multiple minor clonal expansions reflecting various perturbations of the immune homeostasis such as infections, auto-immune diseases, and immunosenescence.

The identification of clones in B-cell populations has been greatly facilitated by the introduction of next-generation sequencing (NGS) techniques which can produce millions of IG sequences simultaneously. The analyse of such sequences can allows the reconstruction of cell lineage, and unravel inter- and intra-clonal repertoire diversity. Several computational methods for clonal grouping have been developed which normally employ some clustering algorithms to infer clonal relationships (*9–11*). For that, they cluster IGH (or IGL) nucleotide (or aminoacid) sequences bearing some similarity. Although these works deal with sequencing data (not with cells), they use the term clone rather than clusters of IGH (or IGL) sequences. For simplicity, we adopted the same terminology.

The performance of clonal grouping methods is hard to evaluate since experimental data containing truly clonally related sequences are difficult to obtain. Most clonal grouping methods are evaluated on simulated repertoires, but most of the time, such data sets might not reflect real cases. It is hard to generate artificial data to simulate all different types of real repertoires, occurring in different physiological or pathological conditions. Thus, the performance evaluation of clonal grouping algorithms requires the development of informative benchmarks (ground truth data). However, generating such data from B-cell repertoires is challenging. Clonally related sequences should have the same V(D)J rearrangement since they derive from a common precursor. Thus, the first step is to predict IGHV, IGHD and IGHJ genes by using some V(D)J assignment tool (*11–14*). Another constraint is related to similarities of CDR3 regions. Sequences with similar CDR3 regions (above a certain threshold) might be members of the same clone. How to determine this threshold is an important point, and for different types of immune responses, the threshold level of mutation indicating clonally related sequences might vary (*5*). Naturally, different clones with different sequence contents will be formed depending on the gene annotation tool and threshold used.

Here we present an automatic method to generate ground truth data from real IGH repertoires. To create benchmarks based on experimental data, we used immune repertoires from individuals with or without malignant lymphoid cells (chronic lymphocytic leukaemia, CLL), but any immune repertoire data can be used. The method addresses the problems mentioned above, because it requires IGHV and IGHJ gene annotations and a fixed CDR3 threshold just to form initial clusters/clones. Next, clusters are refined by allowing sequences to move among different clusters until they find their appropriate place. For that, we optimize two important measures: intra- and inter-clonal diversity. Such measures continually evaluate the consistency of detected clusters/clones until no improvement is observed in their cohesion or separation. By minimizing intra-clonal diversity, we improve the cohesion of each clone, which measures how similar are sequences within the same clone. On the other hand, by maximizing the inter-clonal diversity, we improve the separation among different clones.

## 2 Approach

The method proceeds through two main steps: pre-clustering and refinement. Figure 1 shows its flowchart, and Algorithm 1 the pseudo-code for the refinement step.

**Figure 1:**
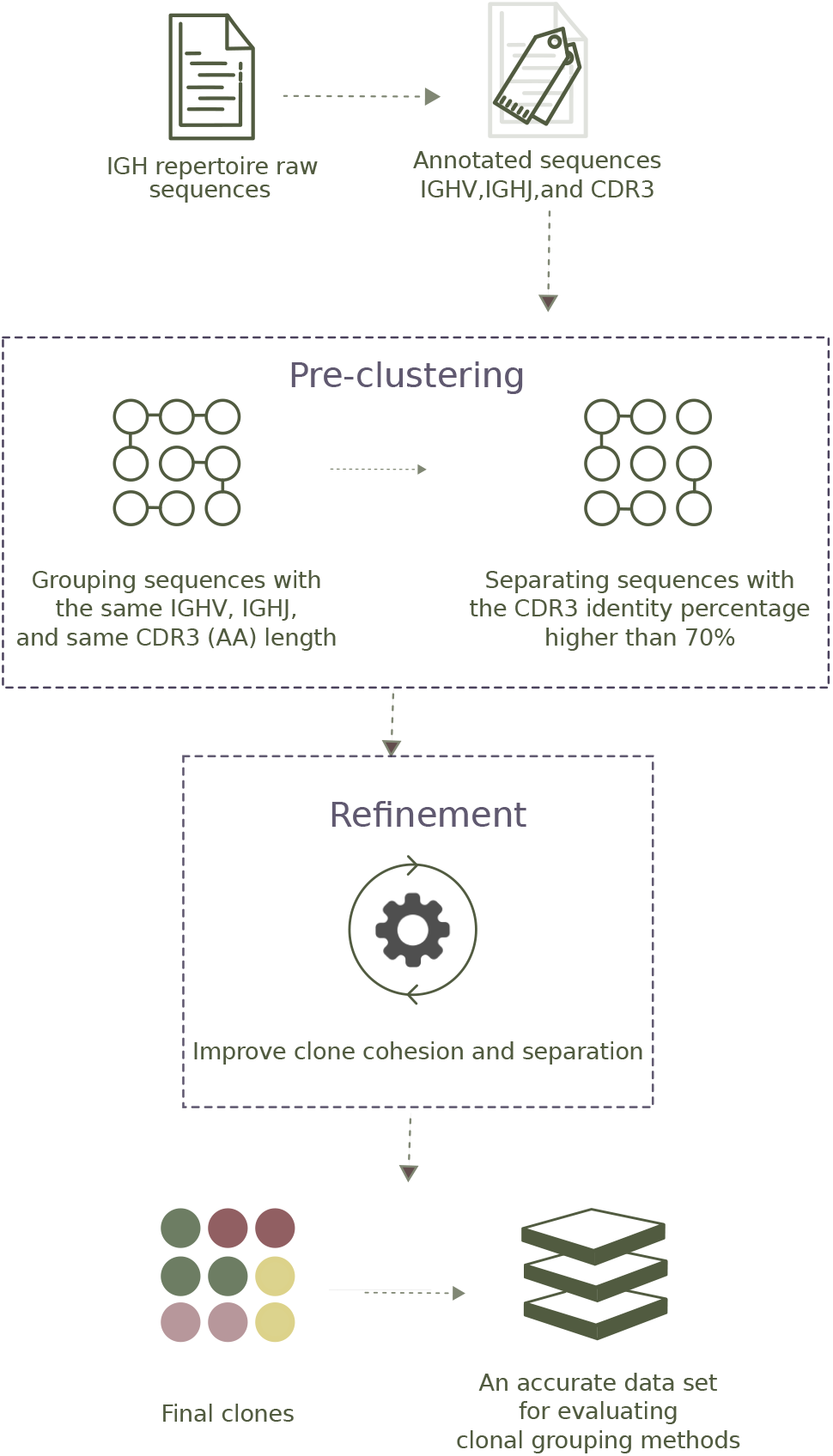
Flowchart of ground truth generation method.

### 2.1 Pre-clustering

The pre-clustering step aims to identify groups of similar sequences to form initial clonal groups that can be refined later. For that, sequences are annotated by IMGT/highv-quest (*12*) to identify their IGHV and IGHJ genes (and alleles), and to locate their CDR3 regions. To form initial clusters/clones, we first group together sequences with the same IGHV and IGHJ genes, and same CDR3 sequence length. Next, we separate sequences with less than 70% of CDR3 identity, see the ‘‘pre-clustering” panel in Figure 1.

### 2.2 Clustering refinement

In this step, we iteratively refine clonal groups until it is impossible to minimize intraclonal distances or maximize interclonal distances. The algorithm described in 1 takes as input the set of initial clusters/clones *C* (generated previously) and for each member sequence *i* ∈ *C* computes two distances: *a_i_* (intraclonal) and *b_i_* (interclonal). Such distances measure the cohesion/separation within detected clusters/clones; they were initially introduced to compute the Silhouette (*15*), a performance measure that helps to interpret and evaluate cluster algorithms when ground truth data are unavailable. In a well-detected cluster, *a_i_* is smaller than *b_i_*, thus, if for a given sequence *a_i_* is higher than *b_i_*, it might indicate that *i* was placed in a wrong cluster, and it should be moved to the cluster with the smallest average distance. If sequences are moved from a cluster *k* to a cluster *l*, then *a_i_* and *b_i_* need be recomputed for all sequences in both clusters. Consequently, each sequence movement launches a new iteration of the algorithm, and it stops if no movement was observed in the previous iteration.

Certainly, the distance metric *d*(*i, j*) (between sequences *i* and *j*) plays an important role when computing *a_i_* and *b_i_*. Distances based on sequence similarity of the whole sequences can be inaccurate since different IGHV/IGHJ genes can present a considerable similarity. Moreover, CDR3 regions are shorter than IGHV/IGHJ genes, and a normalized distance should be more appropriate. Therefore, we split the sequences into three parts, IGHV, IGHJ and CDR3 region, and compute a different distance of each part, separately. The distance *d*(*i, j*) is the arithmetic mean of these three parts and is defined by the equation:

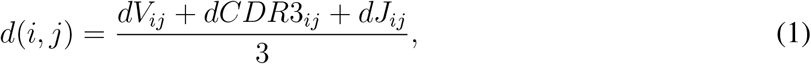

where *dV_ij_* is a binary distance based on IMGT/highv-quest gene identification, it is 0 if *i* and *j* were annotated with the same IGHV gene or 1 otherwise; dCDR3_*ij*_ is the normalized Levenshtein distance (*16*) of CDR3 amino acid sequences; *dJ_ij_* is the normalized Levenshtein distance of IGHJ nucleotide sequences. We recall that the Levenshtein distance computes the minimum number of single-character editions (insertions, deletions or substitutions) required to transform one sequence into the other.

#### Algorithm 1: Clustering refinement

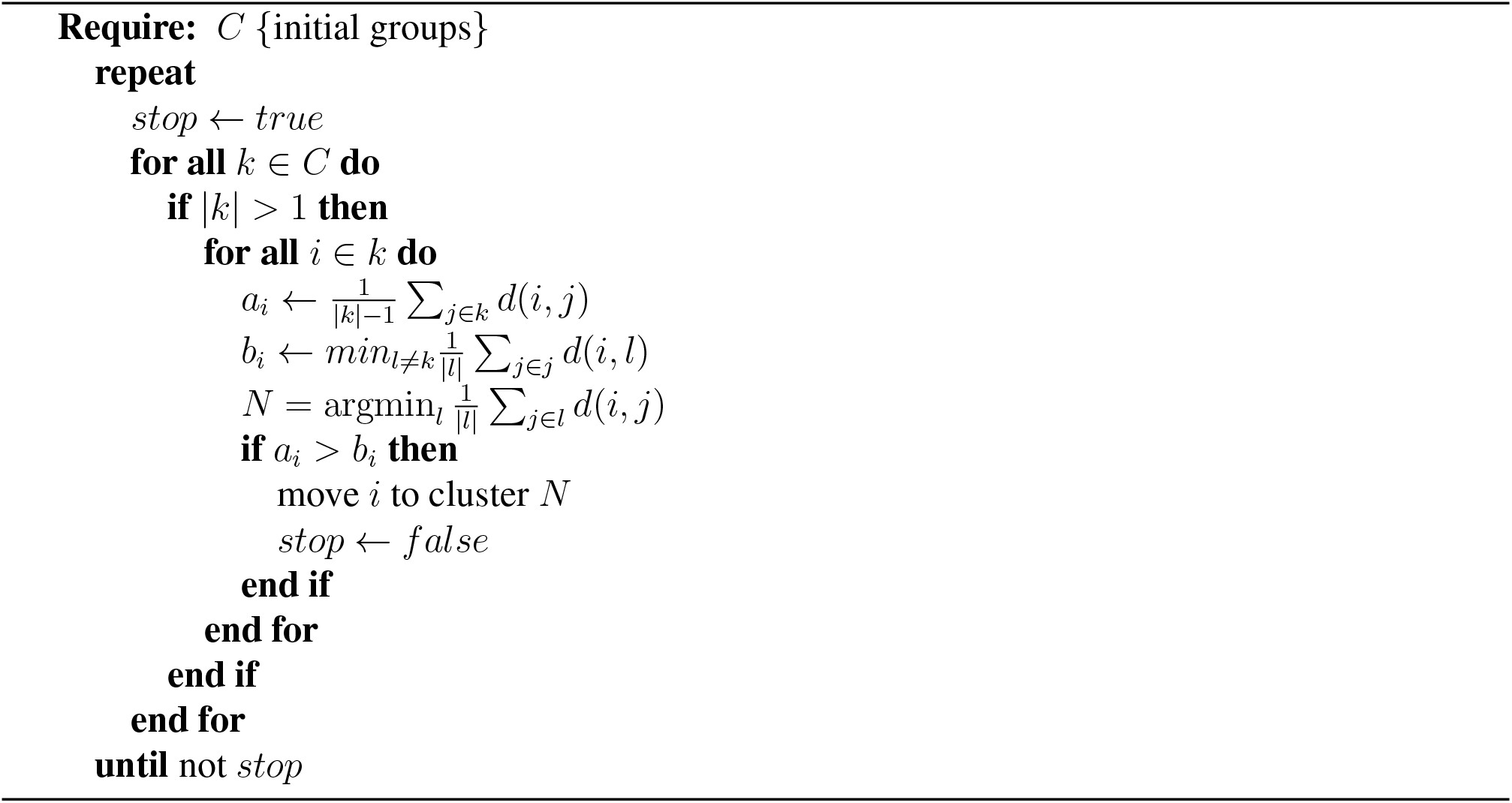

## 3 Methods

### 3.1 Data sets

Two types of ground truth data were generated from artificial and experimental repertoires. Artificial benchmarks were produced by GCTree simulator (*17*), while real benchmarks, based on data from individuals with or without malignant B cells were generated by our method.

#### 3.1.1 Simulated Repertoires

IGH simulated sequences are largely used to evaluate clonal grouping methods (*9,10,18*). Some repertoire simulators have been proposed as part of such tools, but, to the best of our knowledge, an independent B-cell repertoire simulator, that could produce different types of IGH repertoires (clonal, and non-clonal), does not exist. In order to create artificial repertoires, we adapted GCTree (*17*), a B-cell lineage simulator. To generate one repertoire, we ran GCTree several times to produce independent B-cell lineages which together simulate a repertoire.

To produce a B-cell lineage, GCTree randomly selects IGHV, IGHD and IGHJ germline genes from the IMGT database (*19*), then nucleotide(s) can be added to or removed from the IGHV-IGHD and IGHD-IGHJ junction regions. Next, a branching process is performed, and point mutations are included in the descendants. For the branching, GCTree uses an arbitrary offspring distribution that does not require an explicit bounding. Instead, it uses a Poisson distribution with parameter λ to estimate the expected number of offspring of each node in the lineage tree. Somatic hypermutations are simulated by a sequence-dependent process, where mutations are preferentially introduced within certain hot- and cold-spot motifs. GCtree uses the 5-mer context model (*20*) to compute the mutability *μ*_1_,…, *μ_i_*…, *μ_l_* for each residue *i* of a sequence of length *l*. The mutability of the whole sequence *μ*_0_, is then computed by averaging the mutability of its residues: 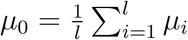. To determine the number of mutations *m* to be introduced in each mutant offspring sequence, GCTree also uses a Poisson distribution with parameter λ_0_, *m* is then computed as *Pois*(*μ*_0_λ_0_); thus, more mutable sequences (higher *μ*_0_) tended to receive more mutations.

GCTree simulator has two main parameters to be set: λ, to estimate the expected number of offspring of each node, and λ_0_, to determine the number of mutations in mutant offspring sequences. We kept λ as the default value (e.g. 2), but we vary λ_0_ to produce simulations with several mutation rates. We have experimented with four values {0.16,0.26,0.36,0.46}, where 0.26 is the default value. Note that higher λ_0_ values produce more divergent B-cell lineages. For each λ_0_ setting, we simulated three types of repertoires: monoclonal, oligoclonal, and polyclonal, obtaining 12 different simulations. The clonal size distribution of each repertoire is shown in Table S1.

#### 3.1.2 Real repertoires

Nine samples of human peripheral blood mononuclear cells collected during routine diagnostic procedures at Pitié-Salpêtrière hospital were selected for this study. Three samples with clonal leukemic cells, and six samples considered as non-clonal (polyclonal) taken from patients devoid of malignancy. Their clonality status had been previously established by conventional methods, including PCR amplification of IGH-VDJ rearrangements followed by Genescan analysis (*21*), see Supplementary Figure S1. DNA sequences were obtained by performing polymerase chain amplification of IGH-VDJ rearrangements followed by NGS paired-end sequencing on an Illumina MiSeq platform. We obtained one “Read 1” and “Read 2” FASTQ files for each sample, which were then merged by the PEAR software (*22*). The merged FASTQ files were converted to FASTA format with seqtk (https://github.com/lh3/seqtk). FASTA sequences were then analyzed using IMGT/High V-QUEST tool (*12*) to identify the IGHV, IGHD and IGHJ genes (and alleles), and delimit the junction and CDR3 regions. The first three columns in Table S2 show the number of reads (sequences), the number of unique sequences, and the clonality status (Supplementary Figure S1) of each repertoire.

### 3.2 Performance evaluation

When ground truth data are available, we can quantitatively assess the ability of clonal grouping algorithms in identifying clonally-related sequences. For that, we apply classical measures such as precision and recall for comparing the inferred clusters/clones to the ground truth data. Consistently, we also compute the F-measure (FM), the harmonic mean of precision and recall, it is an aggregate measure of the inferred cluster’s quality. Precision and recall require three disjoint categories which are: true positive (TP), false-positive (FP) and false negative (FN). Then, we compute precision 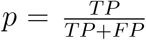, recall 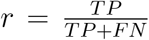, and 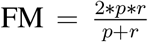. The values of these three metrics are in the interval [0,1], being 1 the best and 0 the worst performance. Certainly, the way TP, FP and FN are computed will affect the accuracy measures. There are at least two ways to compute these values by considering either *pairwise* or *closeness* relationships.

#### 3.2.1 Pairwise

The pairwise procedure considers the binary clustering task, which focus on the relationship between each pair of sequences. A pair of sequences is counted as: TP, if the sequences are found together in both ‘ground truth’ and ‘inferred’ clusters; FP, if the sequences are found separated in the ground truth, but together in the inferred clone; FN, if the pair is found together in the ground truth but separated in the inferred clone, see an illustration in Supplementary Figure S2-A.

#### 3.2.2 Closeness

The closeness procedure first identifies the best correspondence between inferred clusters/clones and ground truth data. It associates cluster pairs that share the maximum of common sequences. Then, for each pair of clusters/clones, saying ‘I’ inferred and ‘G’ ground truth, we compute TP as the intersection between the two sets (*I* ∩ *G*), FP as the difference between inferred and ground truth clusters (*I* \ *G*), and FN as the difference between ground truth and inferred clusters (*G* \ *I*), see an illustration in Supplementary Figure S2-B.

## 4 Results

The goal of clonal grouping is to cluster IGH sequences to facilitate repertoire diversity analysis, detection of potential clonal populations, and lineage reconstruction. Ground truth generation is essential to provide non-trivial data sets that are useful to better evaluate clonal grouping methods, and design practical tools. We have developed a method to automatically generate ground truth data from real IGH repertoires. We first validate it on artificial data where clonal group structures are known. Next, we show how the performance of SCOPe (*10*), a widely used clonal grouping method (*23–25*), varies when using real benchmarks generated by our method. The clonal grouping accuracy was measured by the pairwise and closeness metrics that evaluate the ability to precisely identify clonal memberships, see Section 3.2.

### 4.1 An automatic generation of ground truth data accurately identifies clonally related sequences on simulated repertoires

We first characterized the performance of our ground truth generation method in reconstructing simulated repertoires, where clonal relationships were defined, previously. The goal was to check if one can accurately detect clonal memberships as well as clonal sizes and distributions. Specifically, we created several data sets to simulate several types of repertoires (clonal and non-clonal) with different SHM rates. For that, we adapted GCTree (*17*), a B-cell lineage simulator, which randomly selects germline sequences for generating every lineage and then randomly introduces point mutations in hotspot positions. This process was repeated several times for creating a collection of B-cell lineages, composing a unique repertoire, see Section 3.1.1. To produce simulated repertoire with different SHM loads, we varied the corresponding GCTree parameter λ_0_ that determines the number of mutations in offspring sequences. We experimented four different configurations {0.16, 0.26, 0.36,0.46}, where higher values produce more divergent B-cell lineages. We generated 12 repertoires, totalizing approximately 11k sequences, see initial clonal size distribution in Supplementary Table S1. On these data, we evaluated our method by comparing inferred clusters/clones to truly related clonal sequences generated during the construction of each simulated repertoire.

Our ground truth generation method achieved high precision, recall and F-measure across all simulated data sets for both pairwise and closeness performance measures, Table 1. Across all mutation rates, it was able to accurately identify all pairwise relationships, and reconstruct precisely all repertoires. The absolute performance measures were extremely high, for both clustering accuracy strategies, exhibiting a mean recall over 99%, and a mean precision/F-measure equal to 1. These results show that our method can generate high-quality ground truth data sets.

**Table 1:** Comparing performance of ground truth generation method and SCOPe on simulated repertoires. Third, fourth, and fifth columns show the number of sequences, the number of expected clones, and the number of detected clones, respectively. Pre, Rec, and FM are abbreviation of precision, recall and F-measure, respectively.

| $\lambda_0$ | Clonality | # seq | # exp. clones | # det. clones | Ground truth generation method | | | | | | # det. clones | SCOPE | | | | | |
| --- | --- | --- | --- | --- | --- | --- | --- | --- | --- | --- | --- | --- | --- | --- | --- | --- | --- |
|  |  |  |  |  | Pairwise |  |  | Closeness |  |  |  | Pairwise |  |  | Closeness |  |  |
|  |  |  |  |  | Pre | Rec | FM | Pre | Rec | FM |  | Pre | Rec | FM | Pre | Rec | FM |
| 0.16 | Monoclonal | 958 | 34 | 34 | 1 | 1 | 1 | 1 | 1 | 1 | 35 | 1 | 0.99 | 1 | 1 | 0.58 | 0.73 |
|  | Oligoclonal | 1014 | 43 | 43 | 1 | 1 | 1 | 1 | 1 | 1 | 46 | 1 | 0.97 | 0.99 | 1 | 0.9 | 0.95 |
|  | Polyclonal | 968 | 44 | 44 | 1 | 1 | 1 | 1 | 1 | 1 | 46 | 1 | 0.94 | 0.97 | 1 | 0.88 | 0.94 |
| 0.26 | Monoclonal | 659 | 33 | 33 | 1 | 1 | 1 | 1 | 1 | 1 | 36 | 1 | 0.9 | 0.95 | 1 | 0.6 | 0.75 |
|  | Oligoclonal | 958 | 43 | 43 | 1 | 1 | 1 | 1 | 1 | 1 | 50 | 1 | 0.94 | 0.97 | 1 | 0.65 | 0.79 |
|  | Polyclonal | 964 | 44 | 45 | 1 | 0.99 | 1 | 1 | 0.95 | 0.97 | 49 | 1 | 0.95 | 0.98 | 1 | 0.78 | 0.88 |
| 0.36 | Monoclonal | 924 | 35 | 35 | 1 | 1 | 1 | 1 | 1 | 1 | 36 | 1 | 0.66 | 0.8 | 1 | 0.58 | 0.73 |
|  | Oligoclonal | 991 | 40 | 40 | 1 | 1 | 1 | 1 | 1 | 1 | 57 | 0.99 | 0.66 | 0.8 | 1 | 0.43 | 0.60 |
|  | Polyclonal | 897 | 42 | 43 | 1 | 0.99 | 1 | 1 | 1 | 1 | 58 | 1 | 0.82 | 0.9 | 1 | 0.57 | 0.73 |
| 0.46 | Monoclonal | 952 | 35 | 36 | 1 | 0.99 | 1 | 1 | 0.99 | 1 | 46 | 1 | 0.73 | 0.83 | 1 | 0.16 | 0.28 |
|  | Oligoclonal | 1016 | 43 | 43 | 1 | 1 | 1 | 1 | 1 | 1 | 62 | 1 | 0.84 | 0.92 | 1 | 0.45 | 0.62 |
|  | Polyclonal | 952 | 43 | 43 | 1 | 1 | 1 | 1 | 1 | 1 | 57 | 1 | 0.79 | 0.88 | 1 | 0.57 | 0.73 |

We have also investigated if SCOPe (*10*), a spectral clustering-based method, could produce ground truth data by checking its accuracy on the same 12 simulated repertoires. When considering pairwise performance, SCOPe achieved high precision, recall and F-measure for simulated data sets with lower mutation rates, see Table 1 (rows λ_0_ = {0.16,0.26}). Precision values were equal to 1 for all data sets, while recall and F-measure values were above 0.94 for those six repertoires.

For the remaining data sets, produced with higher mutation rates λ_0_ = {0.36,0.46}, we observed poorer recalls and F-measures. For highly mutated sequences, SCOPe had difficulties in grouping clonally-related sequences, mainly in monoclonal repertoires achieving less than 0.67 of pairwise recall.

SCOPe’s accuracy based on clustering closeness was lower than with pairwise. Closeness evaluation tends to be more challenging than pairwise since clone properties are also evaluated rather than pairwise relationships. As observed with pairwise performance measures, SCOPe obtained high precision values for all data sets, meaning that few false-positives were found in clone grouping. However, lower recall and F-measure values were frequently obtained. We observed a significant difference between the two performance measures overall repertoires, with a significant loss of recall for closeness measures, see Supplementary Figure S3. This difference was even more evident when comparing clonal repertoires. On average SCOPe achieved a pairwise recall of 0.77 vs 0.55 for closeness; pairwise F-measure was 0.92, while closeness was only 0.67. These results highlight SCOPe’ difficulties in reconstructing repertoire populations, showing its limitations as a method to generate ground truth data.

### 4.2 The reconstruction of real repertoires is challenging

So far, clonal grouping methods have been evaluated almost exclusively on simulated repertoires (*9,11,13*). However, clonal sizes, their distributions and the positions of somatic hypermutation of different immune repertoires are hard to simulate. Moreover, such artificial scenarios might not reflect the complexity of real immune repertoires. In order to produce benchmarks that maintain the properties of experimental data, we have applied our algorithm to nine real repertoires from individuals with or without clonal malignant B cells, Section 3.1.2 and Table ??.

Figure 2 shows the clonal distribution for each benchmark, see more details in Supplementary Figures S4-S12. To measure the disequilibrium of a repertoire, we used the Gini index (*26*), which reflects the inequalities among values of a frequency distribution; zero indicates perfect equality, while 1 corresponds to maximal inequality. Clonal repertoires presented the highest Gini index, being close to 1 for individuals 1 to 3. Repertoires 1 and 3 presented similar clonal distributions, with the presence of a major clone representing the quasi-totality of the repertoire, and a small number of minor clones having a low number of sequences, see Figure 2-AC, and Supplementary Figures S4, S6. Individual 2 presented a different clonal distribution with two major clusters, each one taking more than 40% of the repertoire, see Figure 2-B and Supplementary Figure S5. Detailed sequence analysis revealed that the two major clones were composed of a productive and unproductive IGH-VDJ rearrangements, corresponding therefore to a CLL population with biallelic IGH rearrangements, see Supplementary Figures S1-B. Polyclonal cases were split into two groups: (i) those with the predominance of some high-density clusters (*I*_4_, *I*_5_ and *I*_6_), and (ii) the others with a more equilibrated (balanced) sequence distribution (*I*_7_, *I*_8_ and *I*_9_). In the first group, repertoires 4 and 5 presented similar clonal distributions, with Gini index around 0.76 and 0.84, respectively, see Figure 2-DE and Supplementary Figures S7, S8. Individual 6 presented a different configuration from other repertoires in the same group, with the presence of a relatively abundant clone representing 12% of the repertoire, Figure 2-F and Supplementary Figures S9. It is interesting to note that the repertoire 6 has the most biased clonal distribution among polyclonal repertoires with a Gini index of 0.91. In the second group, we found more homogeneous and less biased repertoires; Sample 7 and 8 had similar distributions (Figure 2-GH and Supplementary Figures S10, S11), while repertoire 9 was more irregular (Gini index around 0.83), see Figure 2-I and Supplementary Figure S12. It should be noted however, that in these three cases, the size of the detected clones was small, each of them accounting for less than 1% of the total sequences. Supplementary Table S2 summarizes properties of all benchmarks; we note that the nine benchmarks presented different characteristics, constituting a valuable resource to evaluated clonal grouping methods.

**Figure 2:**
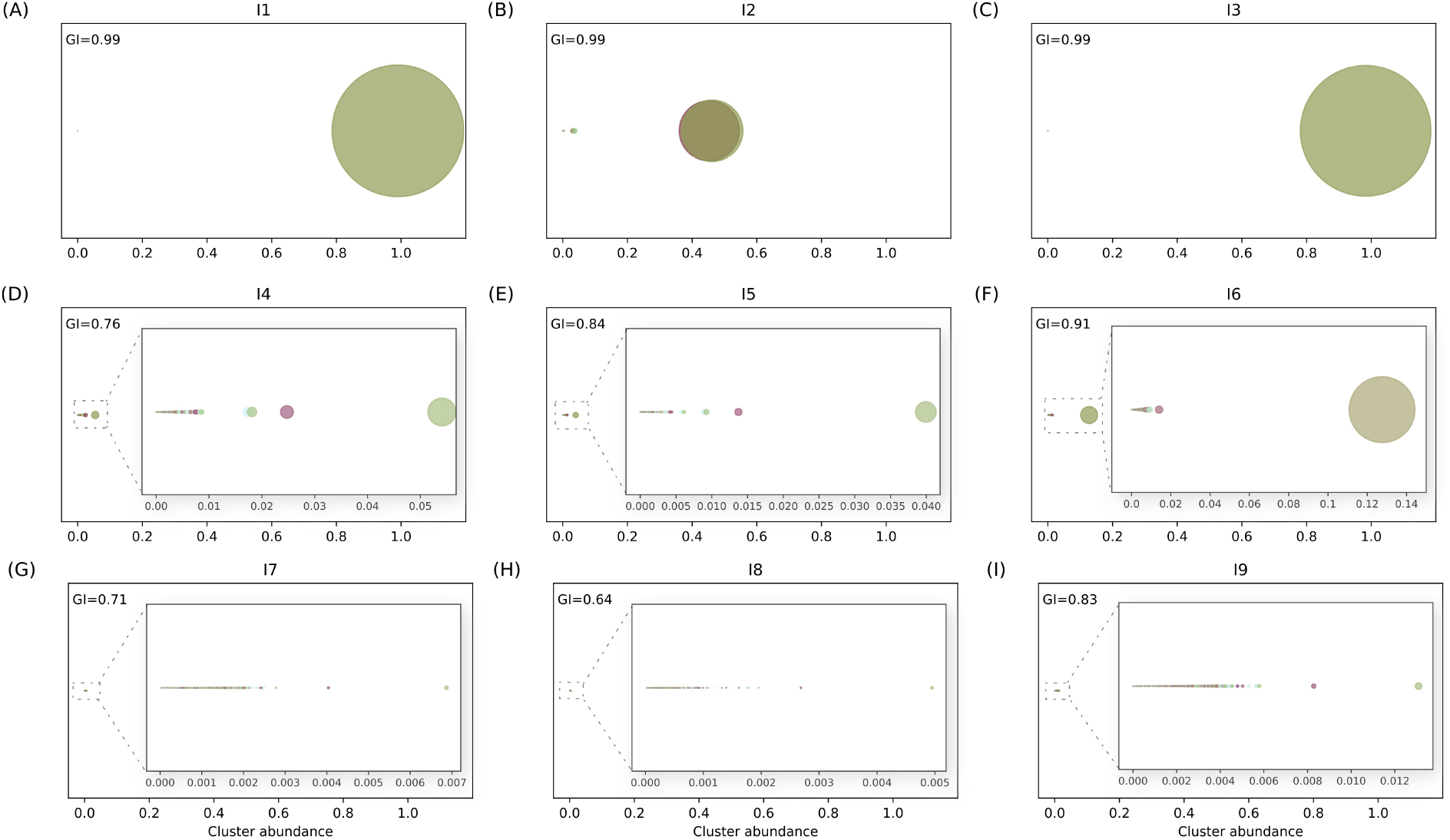
Clonal distribution in real repertoires. Each circle symbolizes a clone, and the clone’s abundance is shown through its size. GI stands for Gini index (*26*) that reflects the inequalities among values of a frequency distribution. To a better visualisation, we zoom samples from I4 to I9.

To demonstrate the issues arising when using experimental data, we ran SCOPe on the nine generated benchmarks and evaluated its performance with two measurements: pairwise and closeness. As pairwise performance can be biased in the presence of highly unbalanced clones, we have evaluated clustering accuracy, gradually. For that, we first ranged detected clones from the most to the less abundant. Then, we evaluated the pairwise performance on the entire set of detected clones. Next, we removed the most abundant clone, and recomputed pairwise performances; we proceeded likewise until all clones were removed.

Table 2 and Supplementary Figure S13 show SCOPe performances on real repertoires. For sample 1, we observed high pairwise performances, but extremely low recall and F-measure closeness, 0.01 and 0.03, respectively. SCOPe was able to detect most of the pairwise relationships in the major clone, but its performance sharply decreased for less abundant clusters/clones, see Supplementary Figure S13-A. Although a slightly better closeness performance was achieved for repertoires 2 and 3, we observed the same tendency as for the repertoire 1, with most of the correct predictions being obtained in clusters with higher density (Supplementary Figure S13-BC), and higher false-positive rates in smaller ones (Supplementary Figure S14-ABC). It is interesting to note that for all clonal repertoires a low F-measure, below 0.2, was obtained for half of the clones, see the black point in Supplementary Figure S14-ABC. In summary, the best closeness performances were achieved in sample 2 and 3, where we observed a less sharp decline in the performance curves when compared to case 1, see Supplementary Figure S13-ABC.

**Table 2:** SCOPe performance on repertoires generated by our ground truth method. Second, third, and fourth columns show the patient identifier, the number of expected clones, and the number of detected clones, respectively. Pre, Rec, and FM are abbreviation of precision, recall and F-measure, respectively. (i) identifies repertoires with the predominance of some high-density cluster/clone, while (ii) identifies more balanced repertoires with no discrepancy between clone sizes.

| Clonality | # ind. | # exp. clones | # det. clones | Pairwise |  |  | Closeness |  |  |
| --- | --- | --- | --- | --- | --- | --- | --- | --- | --- |
|  |  |  |  | Pre | Rec | FM | Pre | Rec | FM |
| clonal | 1 | 162 | 176 | 0.99 | 0.96 | 0.98 | 1 | 0.01 | 0.03 |
|  | 2 | 299 | 89 | 0.98 | 0.97 | 0.98 | 0.99 | 0.29 | 0.45 |
|  | 3 | 324 | 43 | 0.97 | 0.99 | 0.98 | 0.99 | 0.25 | 0.4 |
| non-clonal (i) | 4 | 1770 | 1921 | 0.97 | 0.97 | 0.98 | 0.99 | 0.29 | 0.44 |
|  | 5 | 16226 | 11641 | 0.79 | 0.88 | 0.83 | 0.87 | 0.07 | 0.13 |
|  | 6 | 1975 | 1387 | 0.86 | 0.98 | 0.92 | 0.97 | 0.51 | 0.67 |
| non-clonal (ii) | 7 | 2686 | 2880 | 0.98 | 0.91 | 0.95 | 0.99 | 0.58 | 0.73 |
|  | 8 | 10461 | 7192 | 0.85 | 0.93 | 0.89 | 0.91 | 0.83 | 0.87 |
|  | 9 | 2185 | 2238 | 0.92 | 0.92 | 0.93 | 0.98 | 0.45 | 0.61 |

Among non-clonal individuals of the group (i), we observed similar performances for repertoires 4 and 6, but very different for repertoire 5. Individuals 4 and 6 present different clonal size distributions and different Gini indexes (Figure 2-DF), but they achieved similar pairwise and closeness performance values. We also observed comparable performance curves (Supplementary Figure S13-DF) with slightly better results for case 6. However, SCOPe achieved a lower pairwise precision around 0.86, detecting an important portion of false-positives in the most abundant clone (around 16%), see Supplementary Figure S14-F. On repertoire 5, SCOPe achieved the lowest pairwise performance: 0.79 for precision and 0.88 for recall; it also achieved very low closeness values: 0.07 for recall and F-measure for 0.13. This lower precision indicates a high false-positive rate in the most abundant clone, greater than 10% (Supplementary Figure S14-E). We also notice a faster decreasing in the performance curve of Supplementary Figure S13-E, indicating that good performance is not consistently achieved across all detected clones.

Non-clonal repertoires of group (ii) were more balanced with no discrepancy between clone sizes, see Figure 2-bottom. For individuals 7 and 9, SCOPe achieved high pairwise performances and F-measure closeness greater than 0.6. When evaluating the performance across all detected clones, SCOPe achieved a better performance for the case 7, being the area under the performance curve bigger for case 7, when compared to case 9, see Supplementary Figure S13-G,I. We also noted that the performance curve of the repertoire 9 decreased faster, and half part of clones presented a performance lower than 0.2. For individual 8, SCOPe achieved lower pairwise performance, but the highest closeness values. The low pairwise precision was due to the high false-positive rate of the most abundant clone (around 50%), see Supplementary Figure S14-H; the larger area under performance curve, showed in Supplementary Figure S13-H, could explain the highest closeness performance.

## 5 Discussion

The ability to obtain huge number (millions) of antigen receptor sequences using NGS techniques has dramatically changed our possibilities to explore immune repertoires. Clonally related sequences descend from a common ancestor and present the same V(D)J rearrangement, but they may differ due to the accumulation of somatic hypermutations. Clonal relationships can be computationally identified and IGH most of clonal grouping methods automatically separate sequences into clonal groups based on their similarities, considering either the whole sequence or junction regions. The validation of such methods presents several challenges, since gold standard experimental data, where truly clonal relationships are known with certainty, are difficult to obtain. Simulations can generate ground truth data where the underlying clonal groups are known, but determining the set of parameters to reproduce widely distinct and complex repertoires observed in all types of immune responses is a difficult task. The generation of high precision ground truth data is then an important factor for the development and evaluation of clonal grouping methods, and a high confidence evaluation requires large amounts of data, representing real repertoire clonal distributions.

We have developed a method to automatically generate ground truth data from any IGH repertoire. The method starts by producing initial clusters/clones where sequences with identical IGHV and IGHJ germline annotations, and more than 70% of amino acid similarity on CDR3 regions are considered as clonally related. Next, sequences can move among clusters/clones until a minimum intra-clonal diversity and a maximum inter-clonal diversity are achieved. We validated our method on artificial data that simulated three types of immune repertoires (mono, oligo and polyclonal) with different mutation rates. On the 12 simulated repertoires, the method inferred clonal relationships with very high accuracy, see Table 1, being able to detect clonal memberships and precisely reconstruct repertoire structures. It is interesting to note that on the same set of simulated data, SCOPe, a tool based on spectral clustering, achieved poorer results.

After validation, we generated several benchmarks from real repertoires of individuals with or without clonal B cell populations, the later corresponding to leukemic B cells. We generated nine benchmarks, three from samples with leukemic B cells, and the remaining from non leukemic samples. In all cases, original data structure/distribution was preserved. The data sets presented different properties ranging from very unbalanced clonal distributions to more regular polyclonal repertoires, (Figure 2 and Supplementary Figures S4-S12). To demonstrate that generated benchmarks are challenging data sets, we carried out a comparative analysis of SCOPe performances. For that, we used two different evaluation strategies: pairwise and closeness. The pairwise considers the relationship between all pairs of sequences and has been frequently used for evaluating clonal grouping methods, while the closeness accounts for clone sizes and their distributions. Although closeness performances are often used to evaluate traditional clustering algorithms (*27*), so far they have not been employed for clonal grouping evaluation. On unbalanced datasets both measures have some drawbacks; pairwise tend to bias towards the performance of high-density clusters, hiding the performance of less abundant ones; closeness tends to be very sensitive to changes in the repertoire topology, and overpenalize singleton detection. Therefore, we argue that both strategies should be used to evaluate clonal grouping methods; false-positive rates for each detected clone and performance curves are also necessary.

In contrast to simulated repertoires, SCOPe achieved lower pairwise precision on real samples, obtaining a higher number of false-positives, see mainly repertoires 5, 6, and 8 in Table 2. In real benchmarks, SCOPe had difficulties in grouping clonally related sequences, sometimes mixing sequences from different clusters/clones, and achieving high false-positive rates across all detected clones, see Supplementary Figure S14. Recall/F-measure values were higher for pairwise performance but much lower for closeness. SCOPe performance was not consistent across all detected clones; it achieved better performance for high-density clusters/clones and lower on low-density ones. This is clearly shown in Supplementary Figure S13 with cumulative F-measures from the high to the low-density clusters. SCOPe was unable to reconstruct most of experimental repertoires, achieving lower recalls/F-measures for closeness performance.

In summary, computational methods for grouping IGH sequences into B cell clones are an important part of repertoire studies. The evaluation of such methods is crucial to produce accurate and practical tools. We have implemented a method to automatically generate benchmarks from any experimental repertoire, and we showed how it is important to evaluate clonal grouping with such data sets that preserve repertoire sizes and distributions. We observed very different performances when clonal grouping methods were evaluated on experimental data. These evaluations can contribute to identify problems and give insights to improve the methodology.

