## Supplemental Tables and Figures for "Automatic generation of ground truth data for the evaluation of clonal grouping methods in B-cell populations"

November 23, 2020

### Supplementary Tables

Table 1: **Clonal size distribution for the three types of simulated repertoires.** Each clone is the result of one IGH rearrangement. We only keep the productive simulated sequences; therefore, the final population size might vary from the total sequence count in this table for different simulated datasets..

| Monoclonal | | Oligoclonal | | Polyclonal | |
|---|---|---|---|---|---|
| #Clone | #sequences | #Clone | #sequences | #Clone | #sequences |
| 1 | 700 | 1 | 150 | 14 | 50 |
| 14 | 10 | 1 | 100 | 14 | 10 |
| 12 | 5 | 10 | 50 | 12 | 5 |
| 8 | 3 | 14 | 10 | 8 | 3 |
| 8 | 1 | 12 | 5 | 8 | 1 |
| | | 8 | 3 | | |
| | | 8 | 1 | | |
| 43 | 932 | 54 | 982 | 56 | 982 |

Table 2: **Properties of generated benchmarks**. For each dataset, we show the clonality status, a identifier, the total number of sequences and unique sequences, Gini index, number of total clones and those with at least two sequences

| Clonality | ind | # Sequences | # Unique Sequences | Gini coefficient | # clones | #clones wto singletons | % singletons |
|---|---|---|---|---|---|---|---|
| Monoclonal | 1 | 33599 | 22181 | 0.99 | 162 | 46 | 71 |
| | 2 | 140026 | 84070 | 0.91 | 1975 | 538 | 72 |
| | 3 | 70050 | 61379 | 0.99 | 324 | 146 | 54 |
| Oligoclonal | 4 | 162742 | 104923 | 0.76 | 1770 | 1042 | 41 |
| | 5 | 294203 | 219006 | 0.84 | 16226 | 7818 | 51 |
| | 6 | 57076 | 40673 | 0.99 | 299 | 102 | 65 |
| Polyclonal | 7 | 73888 | 53093 | 0.71 | 2686 | 1321 | 50 |
| | 8 | 61990 | 19949 | 0.64 | 10461 | 5241 | 49 |
| | 9 | 65853 | 26431 | 0.83 | 2185 | 869 | 60 |

### Supplementary Figures

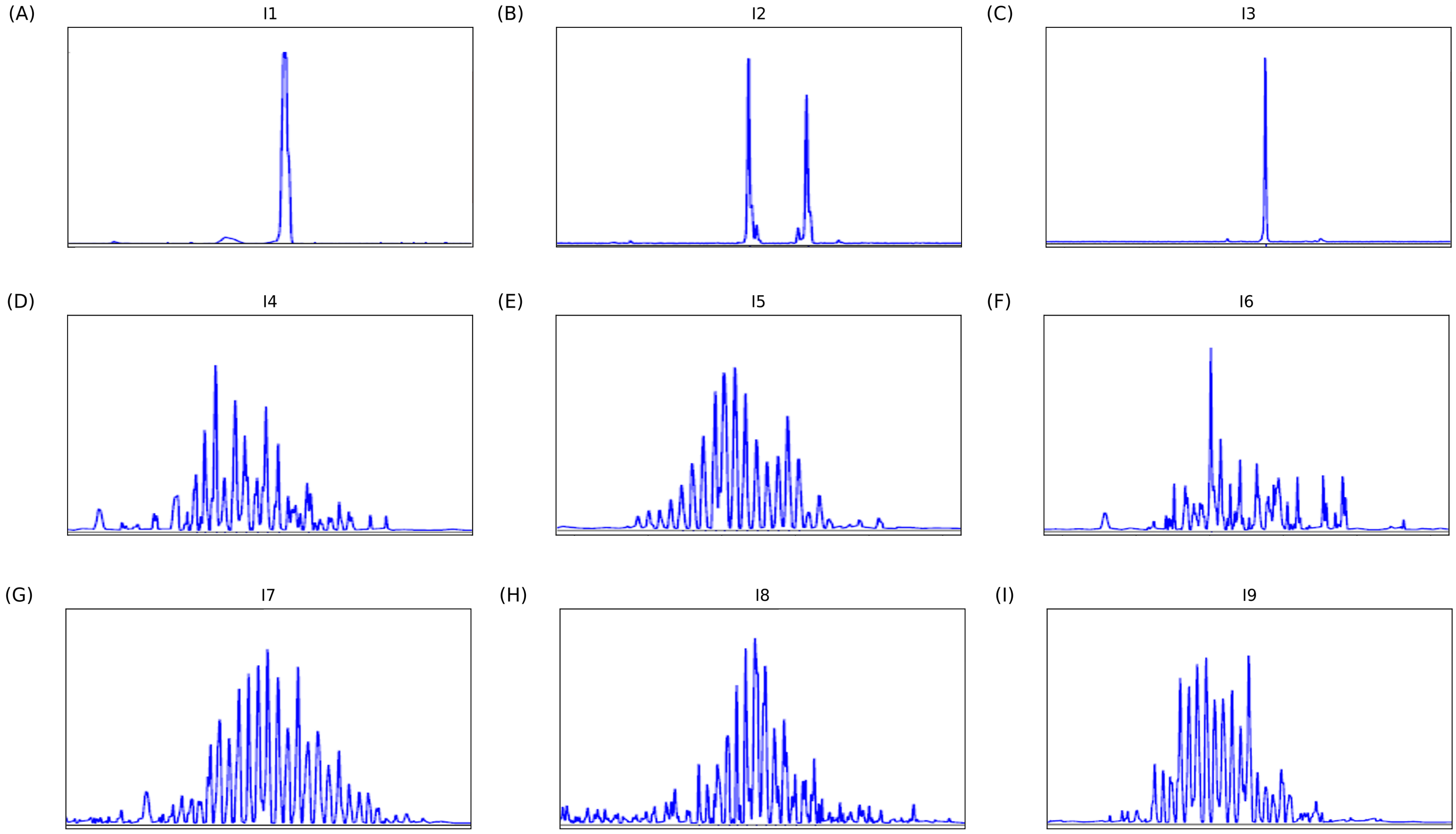


Figure 1: **GeneScan profiles of samples** IGH-VDJ rearrangements were amplified using conventional methods and PCR products further analyzed by capillary electrophoresis. (A-C) Samples from individuals with monoconal B-cell malignancy : monoallelic profile (A and C) or biallelic profile (B); (D-I) non-malignant samples : regular polycylonal profile (D, E, G, H, I) or irregular polyclonal profile (F).

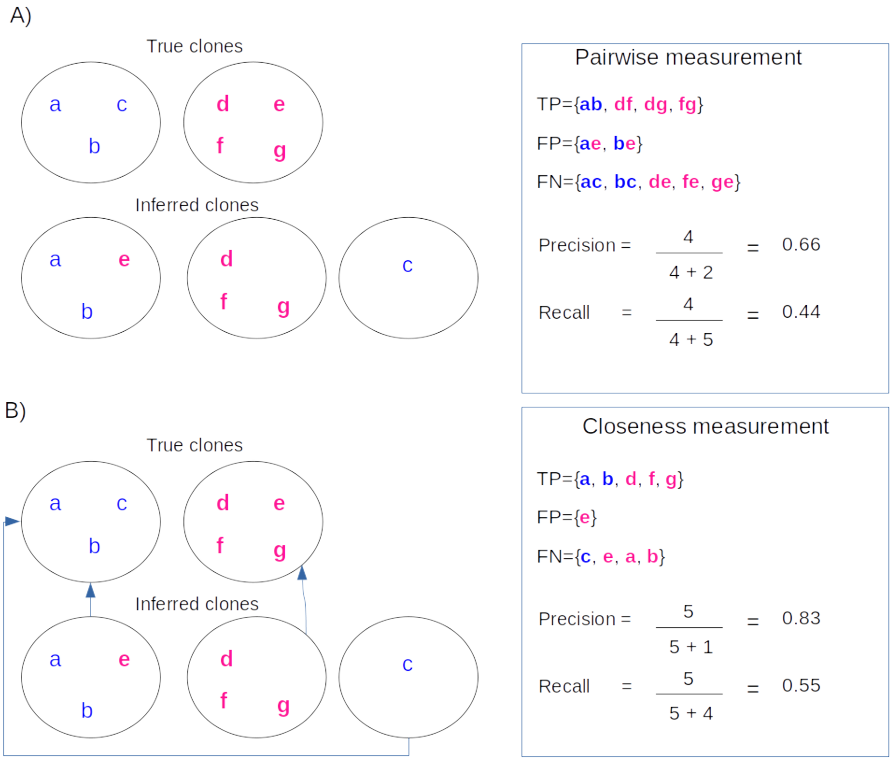


Figure 2: **Clustering performance measures**

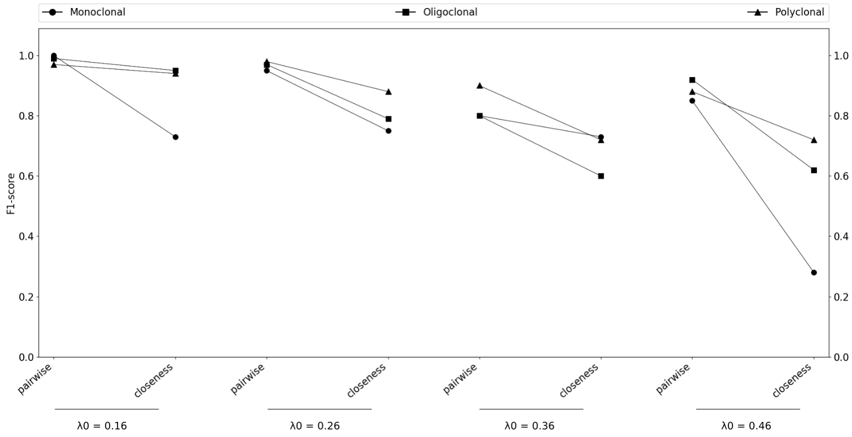


Figure 3: **Comparison of SCOPe performances on simulated repertoires**. For each mutation rate, we compare pairwise and closeness measures in the three type of repertoires: monoclonal, oligoclonal and polyclonal

**Legend for Figures 4 to 12**

A) Circle representation of the clone abundance. Each circle symbolizes a clone, and the clone's abundance is shown through its size.

B) Number of sequences in each clone, all clones are represented, vertical axe is in log scale.

C) Lorenz curve and Gini coefficient. A Lorenz curve shows the graphical representation of clonal inequality. On the horizontal axe, it plots the cumulative fraction of total clones when ordered from the less to the most abundant; On the vertical axe,it show the cumulative fraction of sequences.

D) Percentage of the 100 most abundant clone

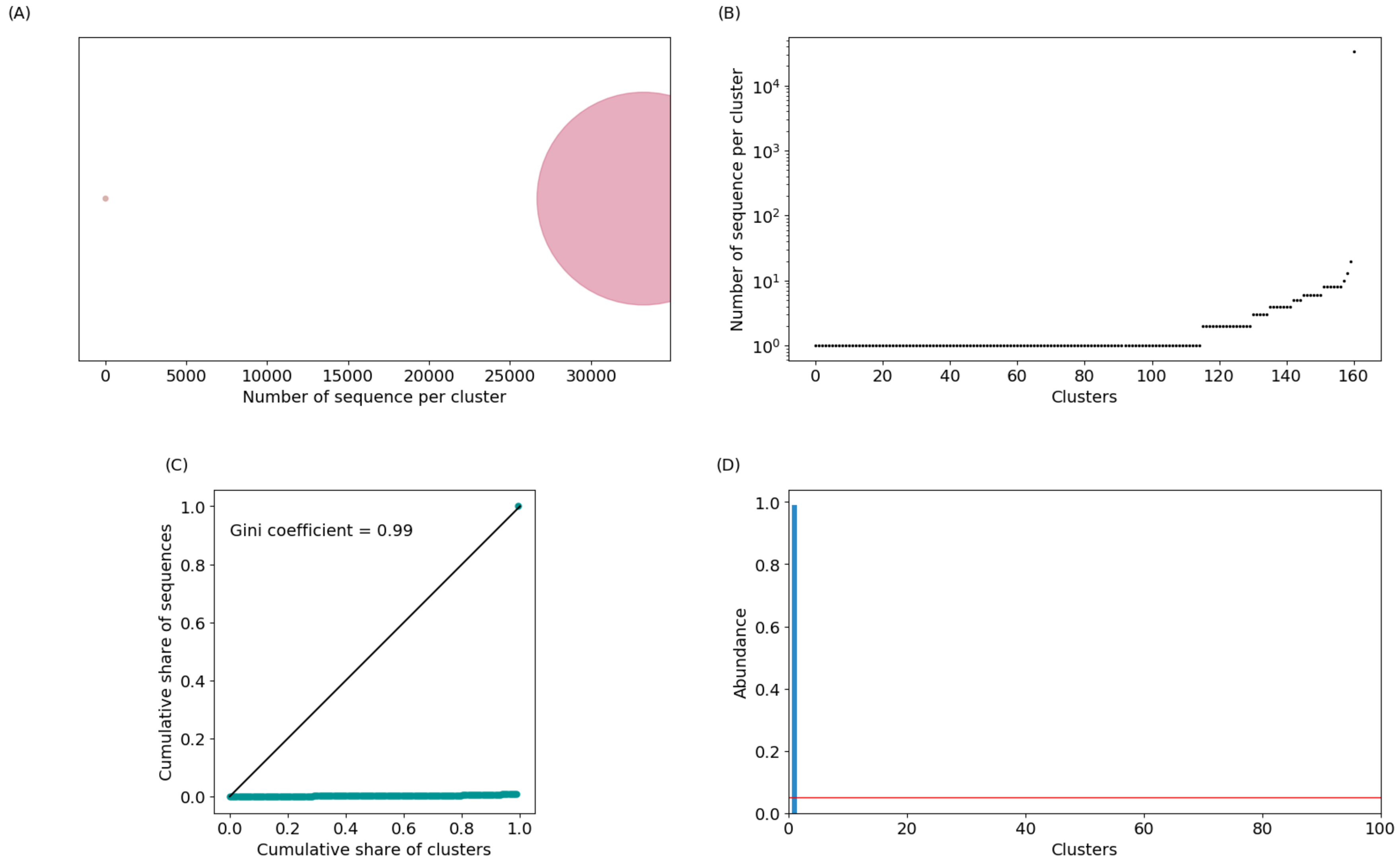


Figure 4: **Repertoire of individual 1**

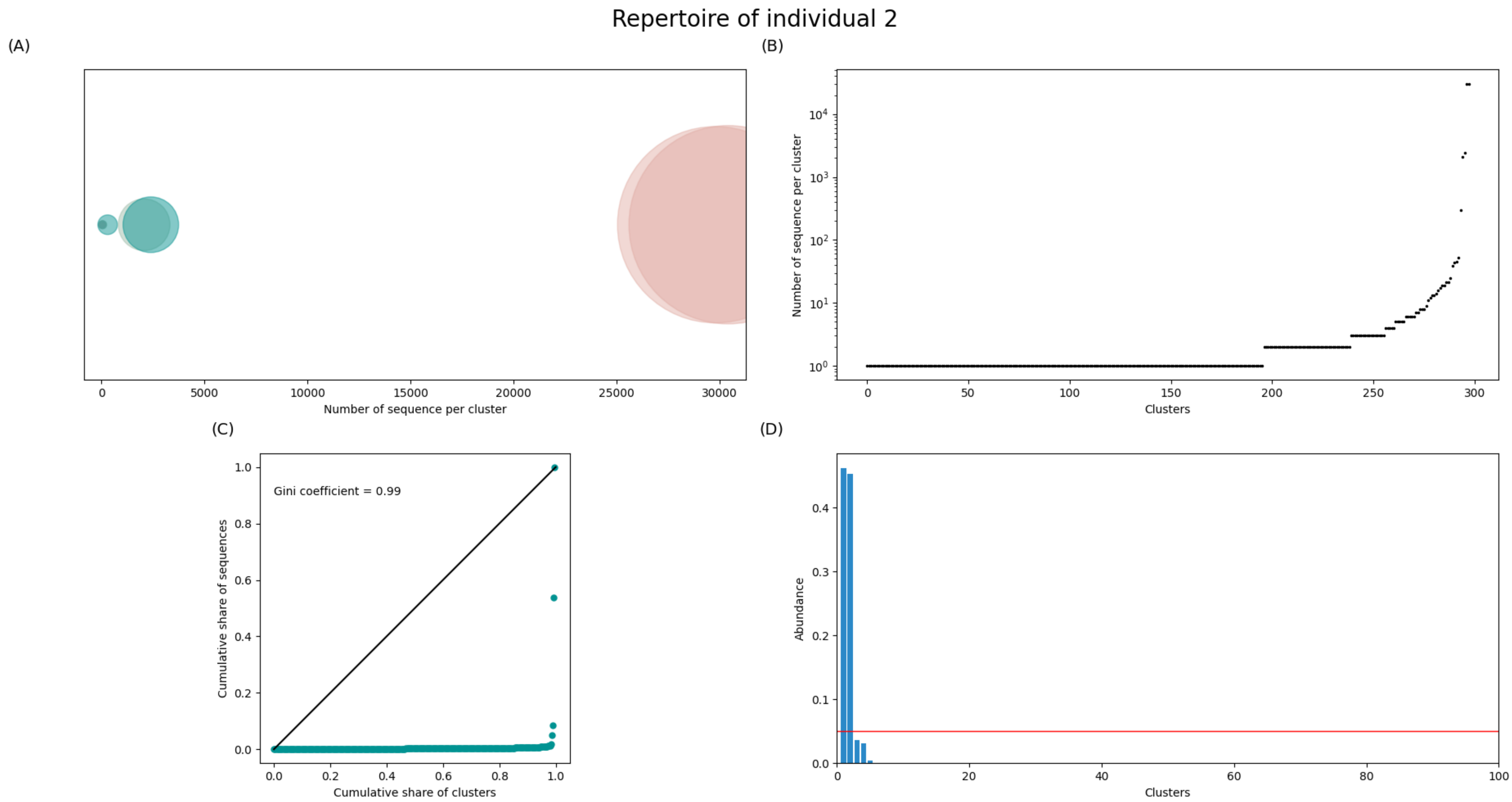


Figure 5: **Repertoire of individual 2**

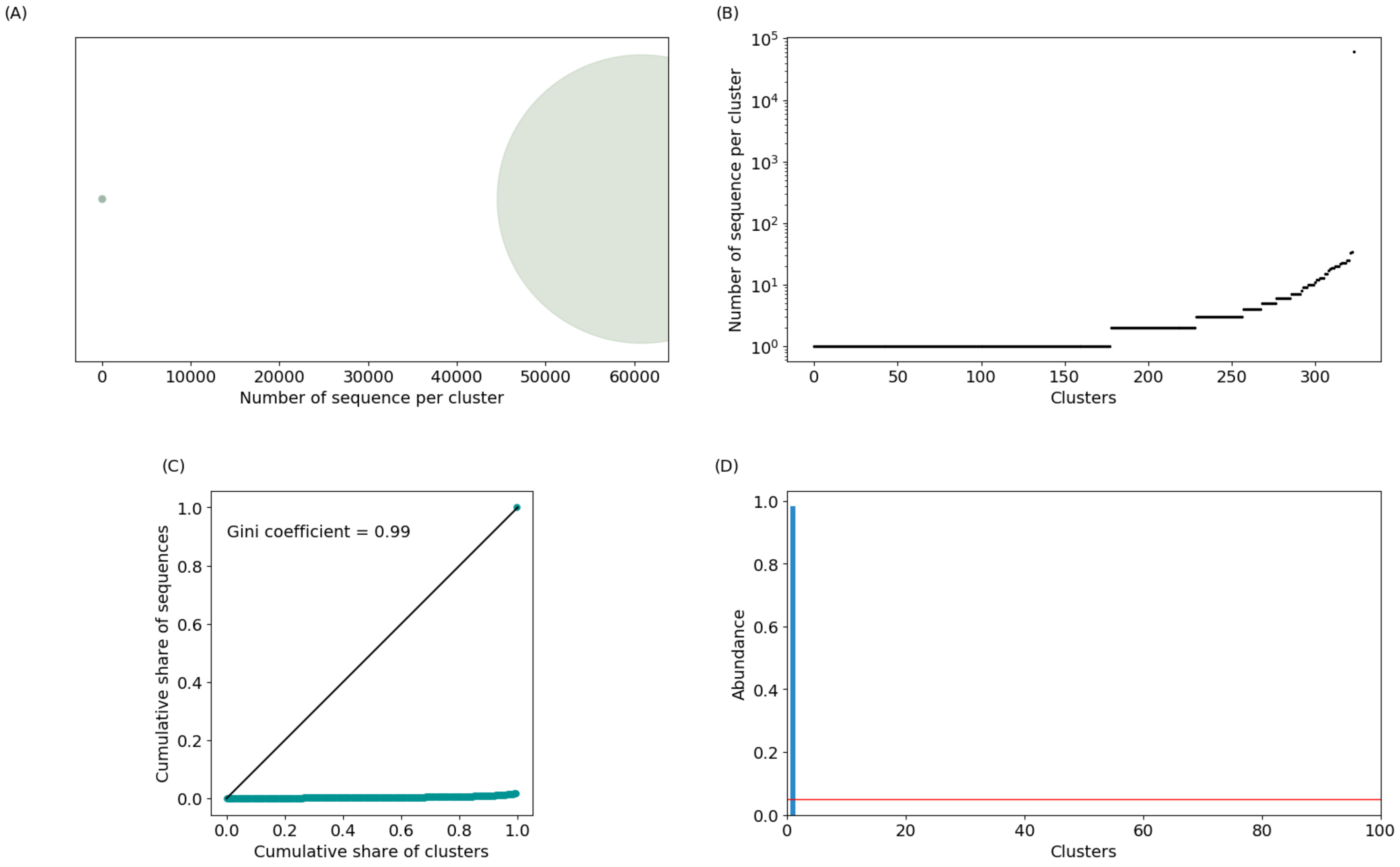


Figure 6: **Repertoire of individual 3**

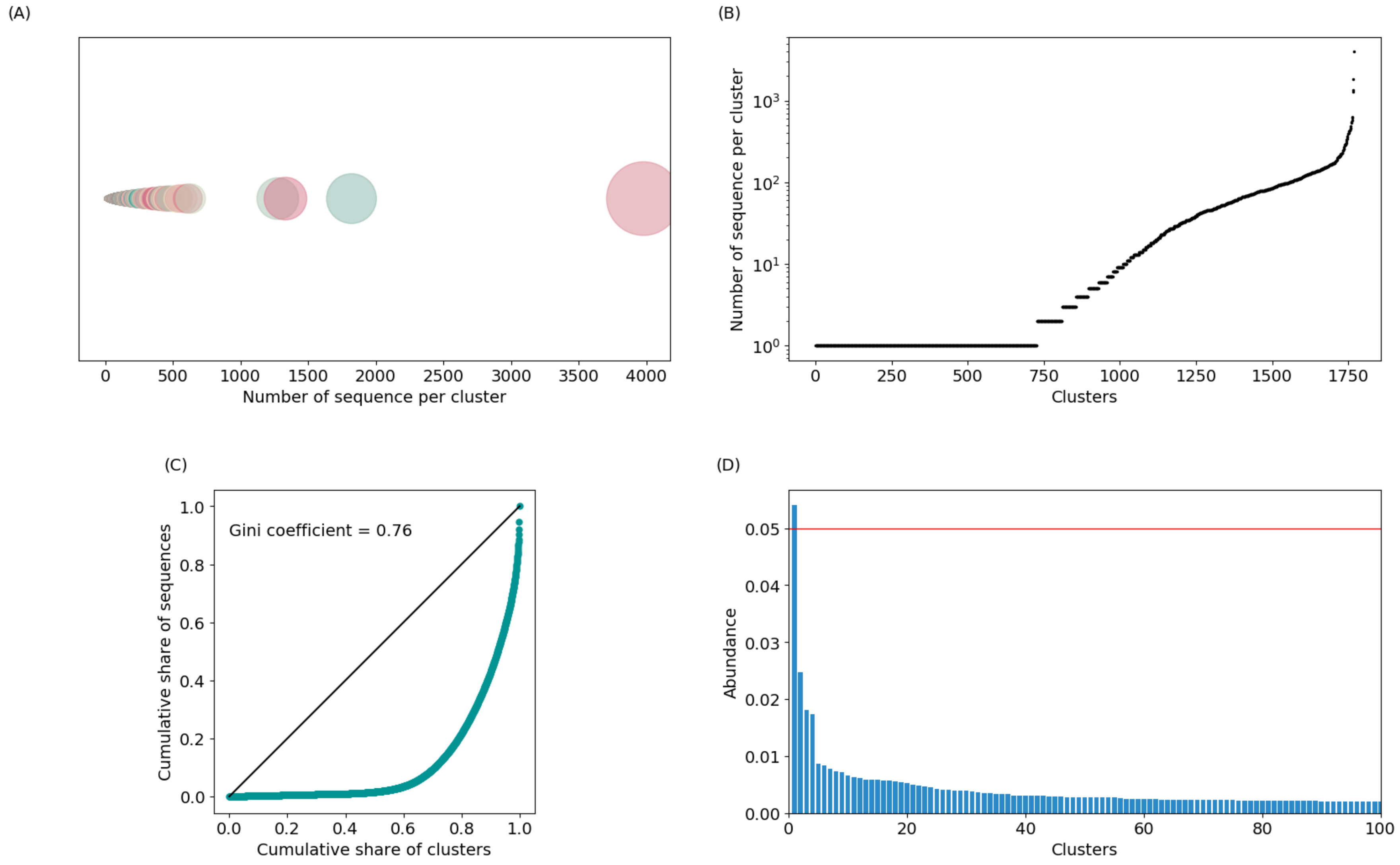


Figure 7: **Repertoire of individual 4**

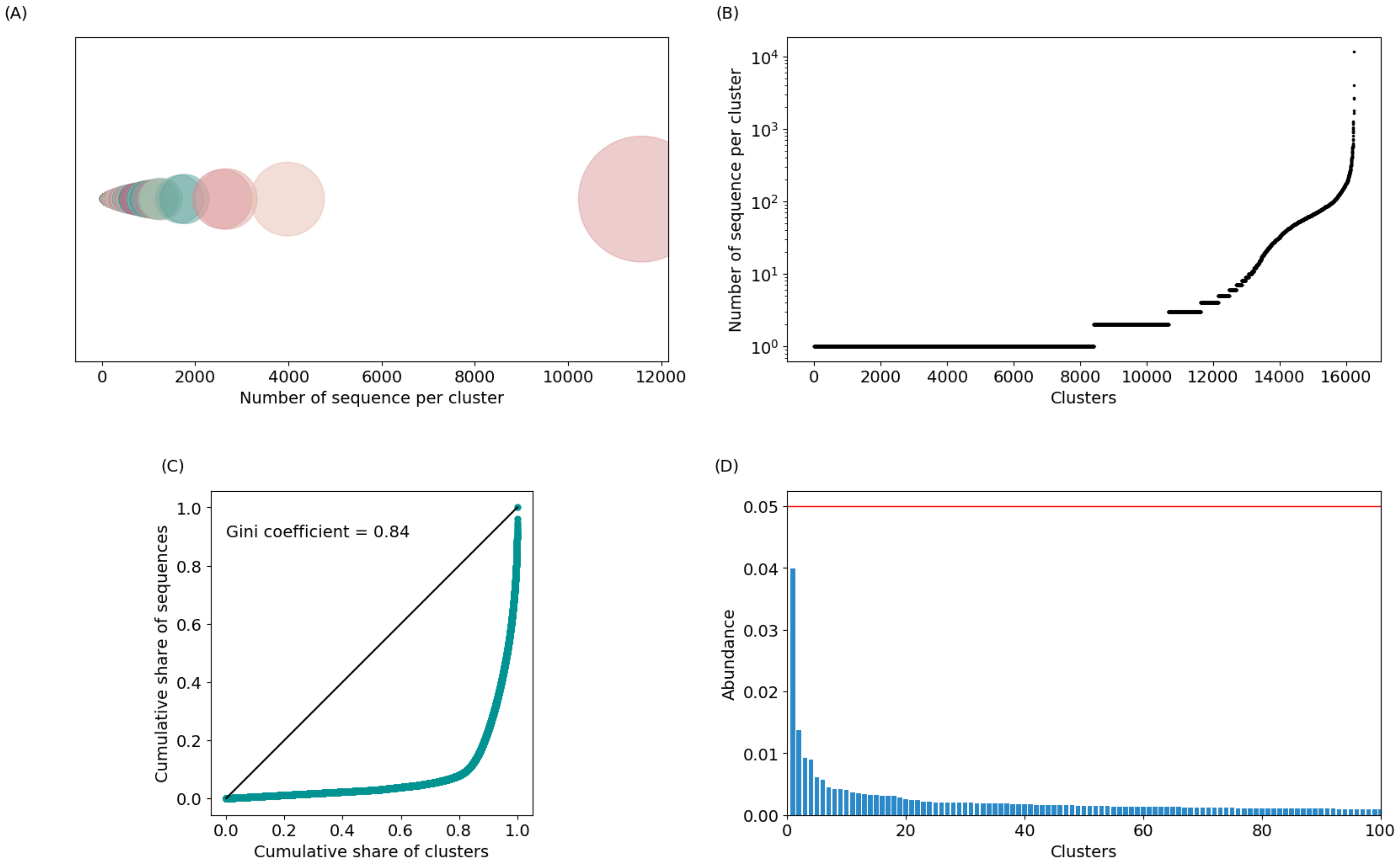


Figure 8: **Repertoire of individual 5**

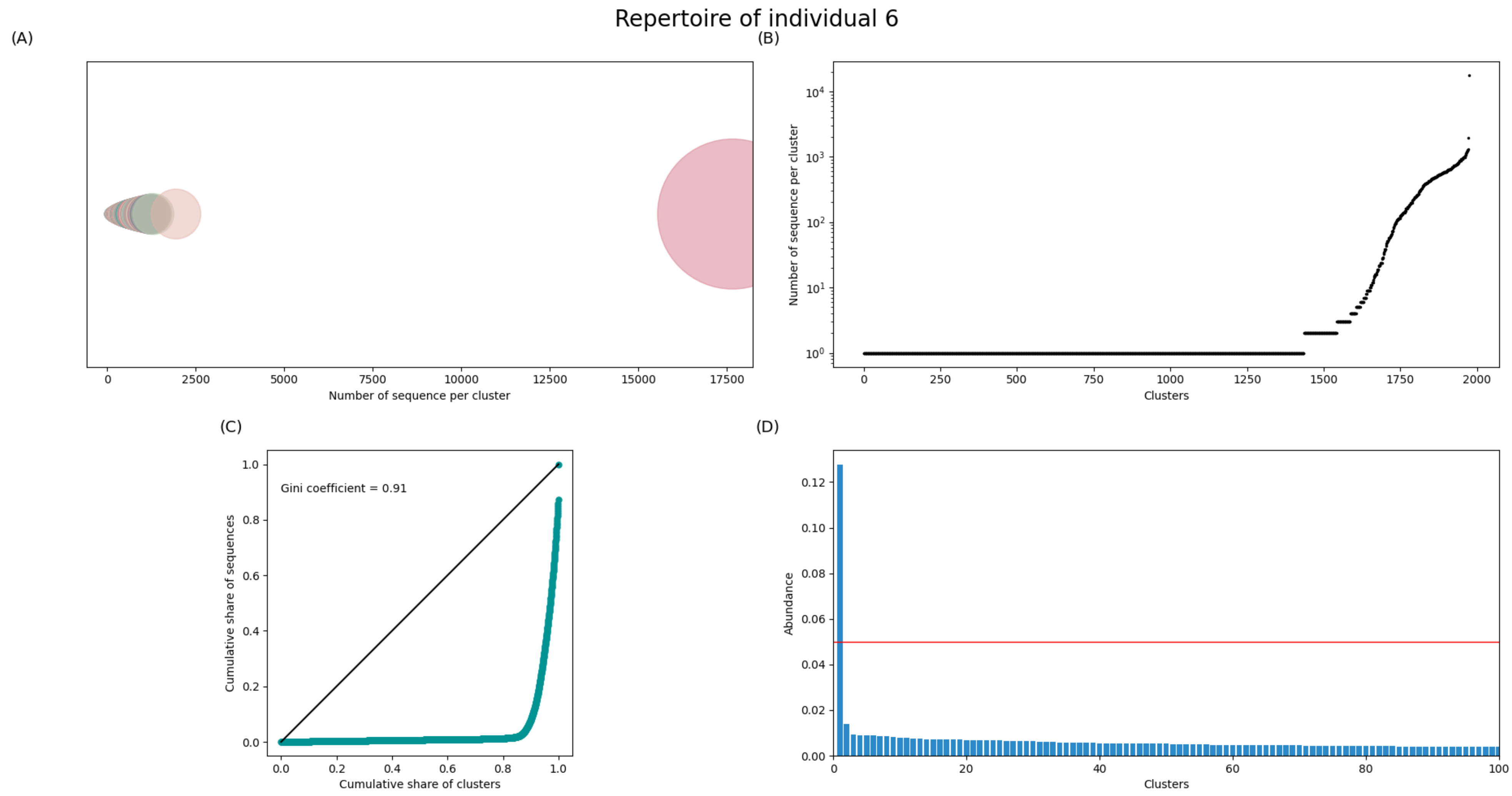


Figure 9: **Repertoire of individual 6**

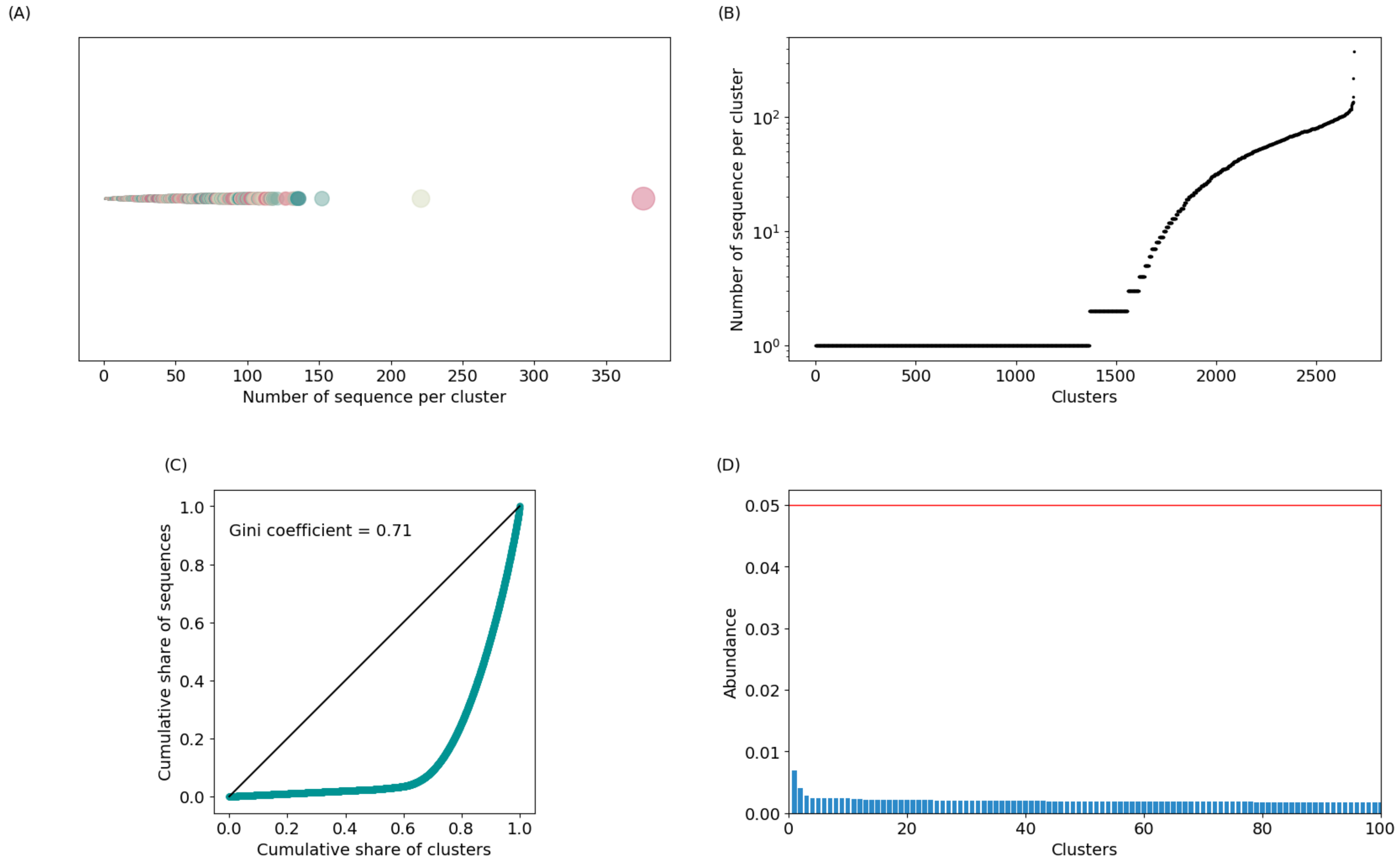


Figure 10: **Repertoire of individual 7**

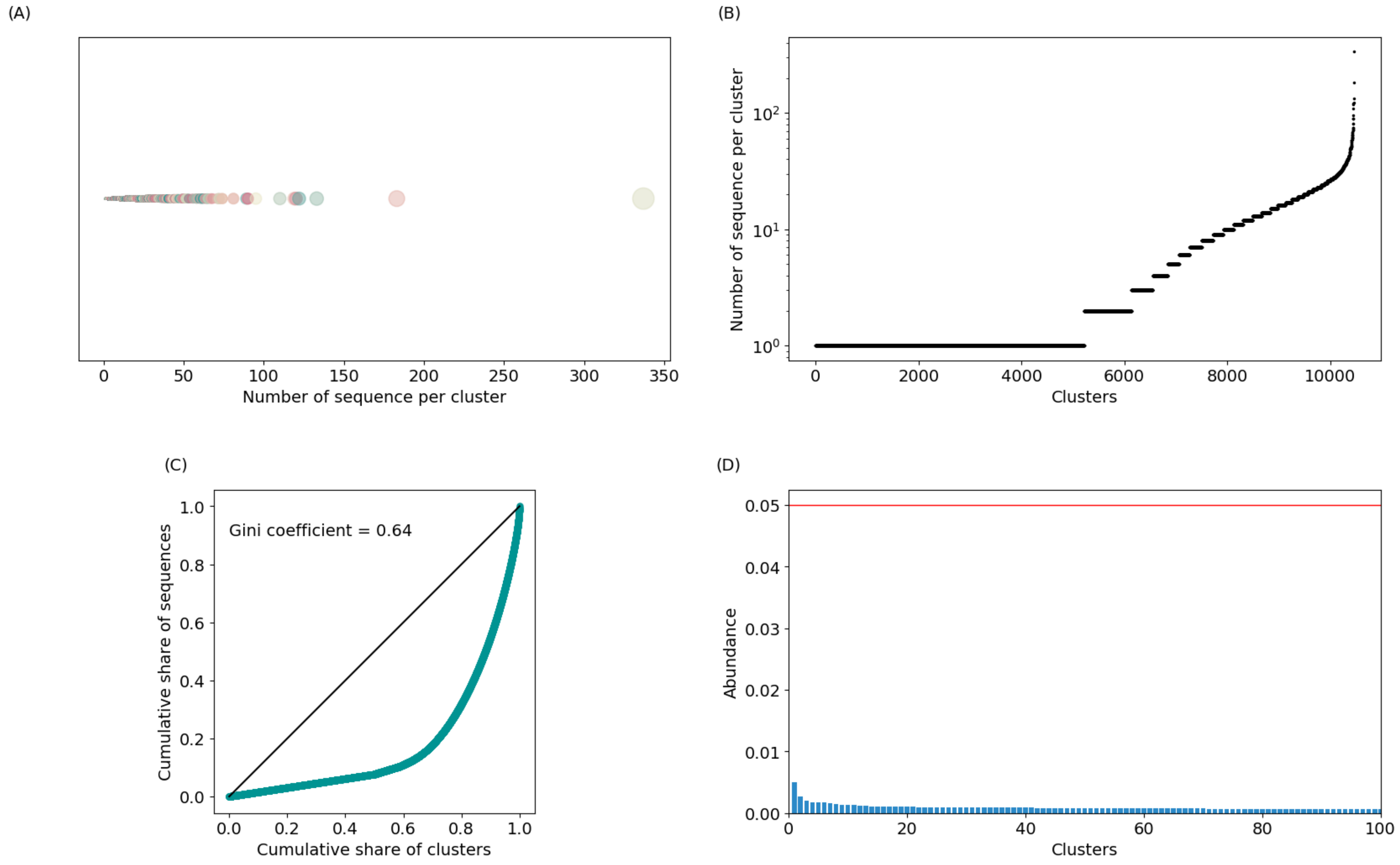


Figure 11: **Repertoire of individual 8**

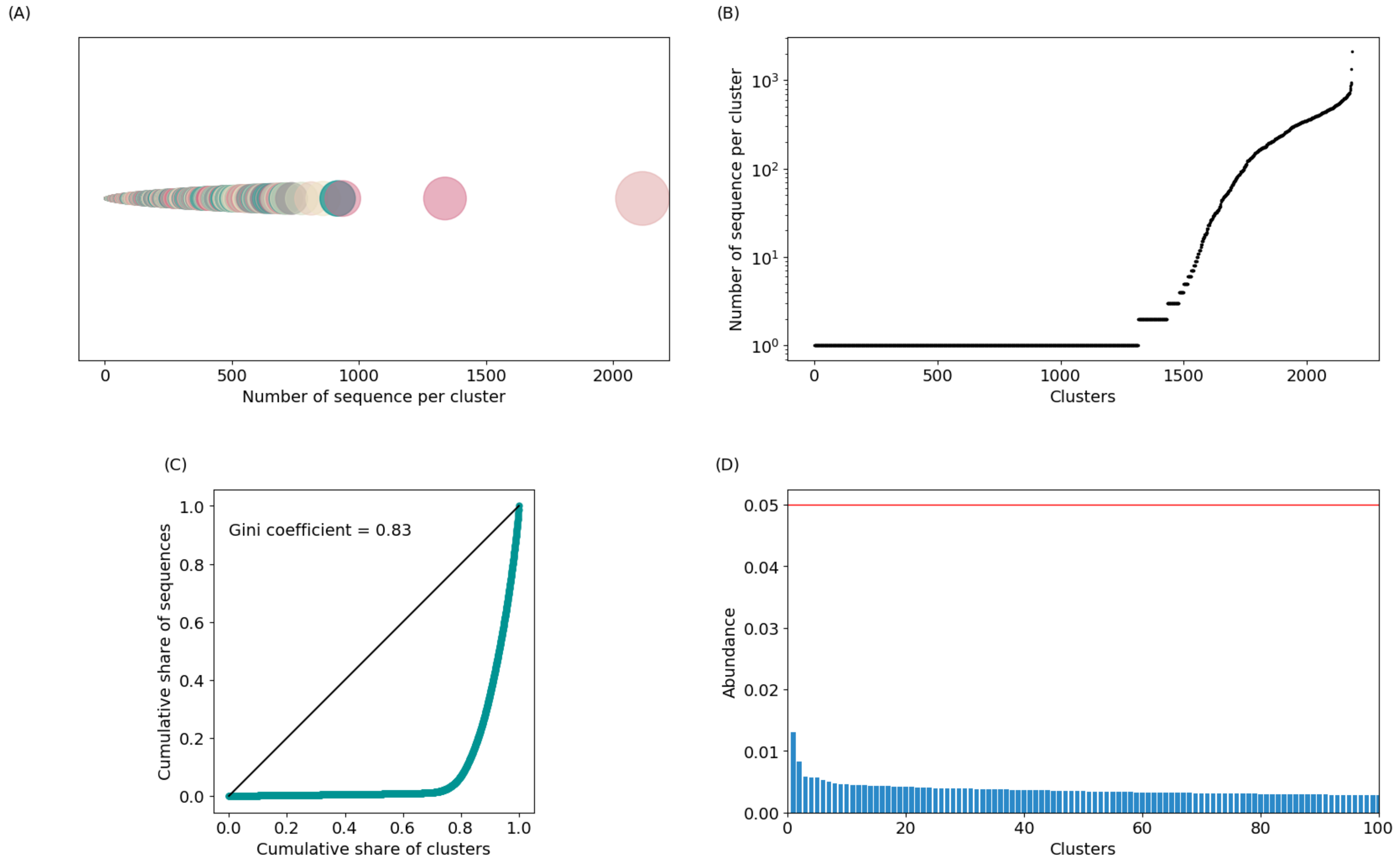


Figure 12: **Repertoire of individual 9**

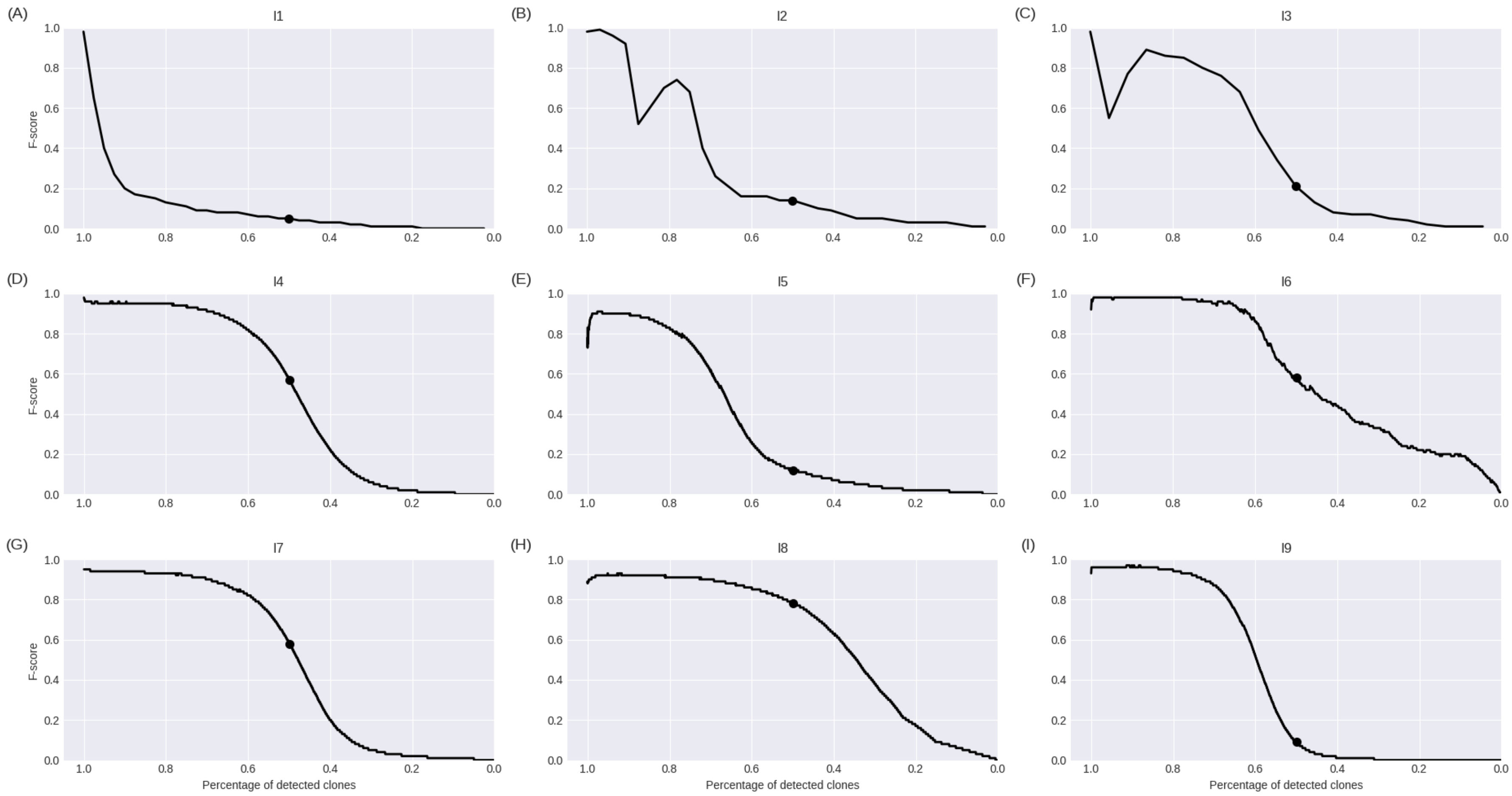


Figure 13: **Performance curve of SCOPe clonal grouping**

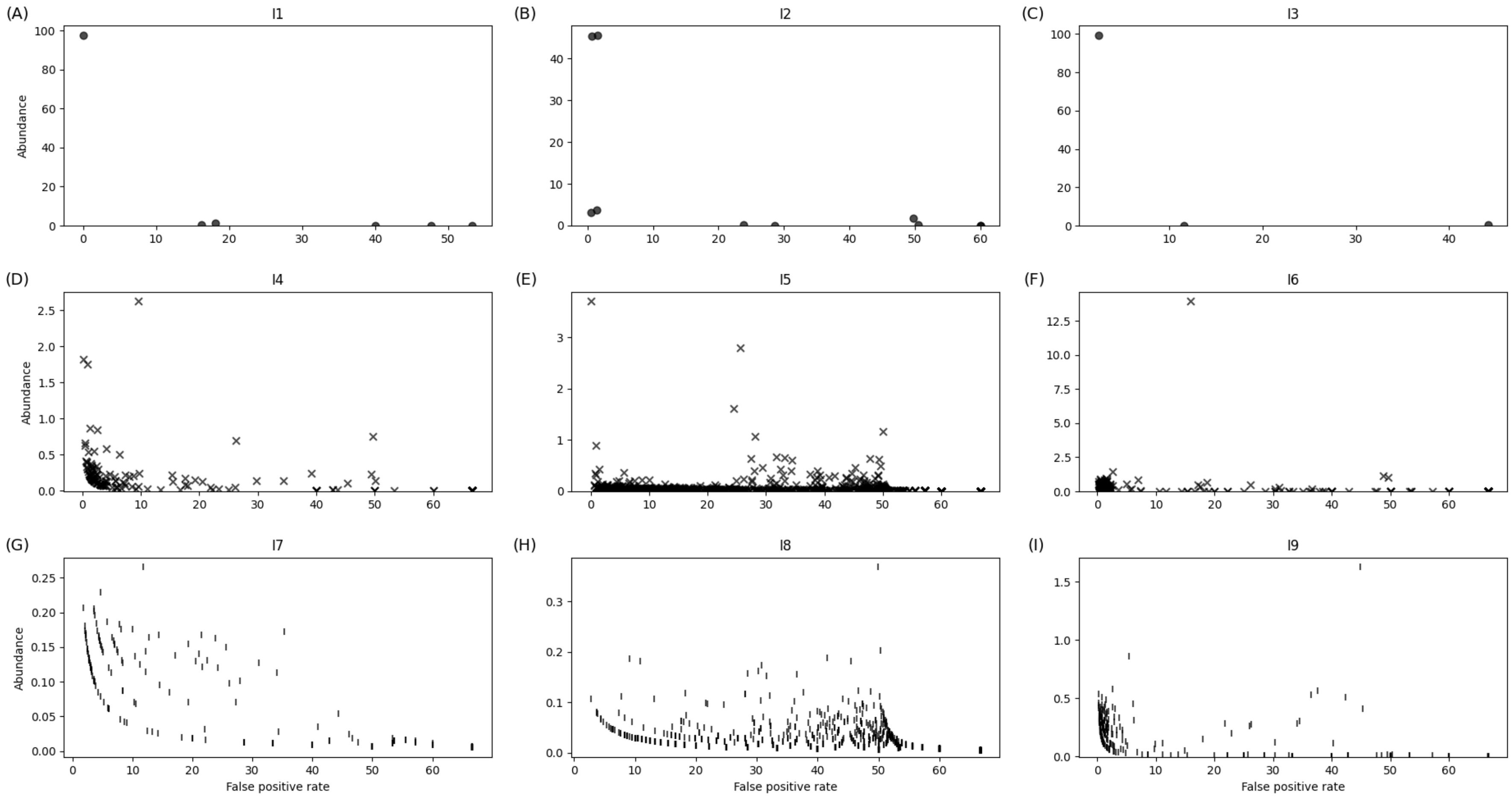


Figure 14: **False-positive rates of SCOPe clonal grouping**
